# A DEAD-box helicase drives the partitioning of a pro-differentiation NAB protein into nuclear foci

**DOI:** 10.1101/2022.03.28.486141

**Authors:** Akiko Doi, Gianmarco D. Suarez, Rita Droste, H. Robert Horvitz

**Affiliations:** Howard Hughes Medical Institute, Department of Biology, Massachusetts Institute of Technology, Cambridge, MA 02139, USA; The Hong Kong University of Science and Technology, Clear Water Bay, Kowloon, Hong Kong

## Abstract

How cells regulate gene expression in a precise spatiotemporal manner during organismal development is a fundamental question in biology. Recent studies have demonstrated the role of transcriptional condensates in gene regulation^1–5^. However, little is known about the function and regulation of these transcriptional condensates in the context of animal development and physiology. We found that the evolutionarily conserved DEAD-box helicase DDX-23 controls stem cell fate in *C. elegans* at least in part by binding to and facilitating the condensation of MAB-10, the *C. elegans* homolog of mammalian NAB protein. MAB-10 is a transcriptional cofactor that functions with the EGR protein LIN-29 to regulate the transcription of genes required for exiting the cell cycle, terminal differentiation, and the larval-to-adult transition^6^. We suggest that DEAD-box helicase proteins function more generally during animal development to control the condensation of NAB proteins important in cell-fate decisions and that this mechanism is evolutionarily conserved. In mammals, a comparable mechanism might underlie terminal cell differentiation and when misregulated might promote cancerous growth.

## Main

During animal development, gene regulation must be temporally precise for proper cell-fate decisions to occur. A key evolutionarily conserved pathway that governs this process consists of developmental timing (heterochronic) genes. Heterochronic genes were first discovered in *C. elegans* from genetic screens for mutants with altered developmental timing of the stem cell-like population of hypodermal seam cells^7^. The *C. elegans* heterochronic genes execute a temporally regulated larval developmental program that drives a precise succession of events, culminating in the terminal differentiation of the seam cells and the onset of adulthood^8–11^. The mammalian homologs of these genes play critical roles in various aspects of physiology and development including metabolism, stem-cell regulation, and fertility, and dysregulation of heterochronic genes has been implicated in many cancers^12–19^.

During *C. elegans* development, the heterochronic gene *lin-28* is expressed during the early larval stages and its protein product, LIN-28, inhibits the processing of the *let-7* microRNA^7, 10, 14, 20^. In the final larval stage, *let-7* microRNA levels are up-regulated, silencing LIN-41 and thereby relieving *lin-29a* repression by LIN-41^21–23^. LIN-28 also acts independently of *let-*7 to down-regulate *lin-29b* levels via the *hbl-1* gene^23^. These coordinated LIN-28 activities inhibit LIN-29, the central driver of exit from the (stem) cell cycle, terminal differentiation and the larval-to-adult transition^21, 24^. LIN-29 is an <u>e</u>arly <u>g</u>rowth response (EGR) protein that acts in part with the <u>N</u>GFI-<u>A</u>-<u>b</u>inding protein (NAB) transcriptional co-factor MAB-10 to control both the expression of genes that regulate terminal differentiation in the hypoderm and the onset of adulthood^6^. This process is evolutionarily conserved. For example, in mammals EGR proteins interact with NAB proteins to regulate terminal differentiation in multiple lineages^25, 26^ and the onset of puberty^17, 27^. Furthermore, EGR1, the mammalian homolog of LIN-29, is a barrier to the reprogramming of somatic cells into human induced pluripotent stem cells (iPSCs)^16^, highlighting the key involvement of this gene in controlling stem-cell fates. Despite the importance of this pathway, mechanisms that regulate the transcriptional activities of the terminal effectors LIN-29 (EGR) and MAB-10 (NAB) remain largely unknown.

To better understand mechanisms that control the activity of the *C. elegans* developmental timing NAB protein MAB-10, we analyzed MAB-10 genetically and biochemically. Here we report that the DEAD-box helicase gene *ddx-23* is a regulator of stem-cell fate and that DDX-23 binds to the MAB-10 protein and facilitates the formation of MAB-10 nuclear foci. We propose that these MAB-10 structures control the transcriptional landscape in terminally differentiated cells.

We further propose that the interaction between DDX-23 and MAB-10 is evolutionarily conserved, that a similar mechanism regulates mammalian stem-cell development and the onset of puberty, and that misregulation of this process can promote cancer progression.

### MAB-10 protein forms nuclear foci *in vivo*

To better define the role of MAB-10 during *C. elegans* development, we characterized the MAB-10 expression pattern and MAB-10 protein dynamics. To examine MAB-10 expression and localization under native regulatory control, we endogenously tagged the C-terminus of the *mab-10* genomic locus with GFP using the CRISPR-Cas9 genome editing technology. The MAB-10::GFP animals did not display any of the phenotypic abnormalities characteristic of *mab-10(lf)* mutants, indicating that this GFP-tagged *mab-10* locus maintains *mab-10* function. MAB-10::GFP expression turned on in the nuclei of two classes of epithelial-like cells -- hypodermal cells and seam cells -- in the early L4 stage. As the animals completed the larval-to-adult transition and became young adults, MAB-10::GFP expression increased and became localized throughout nuclei as well as in nuclear foci. In one-day old adults, most of the MAB-10::GFP signal was localized to nuclear foci (**Fig. 1a**).

**Figure 1.**
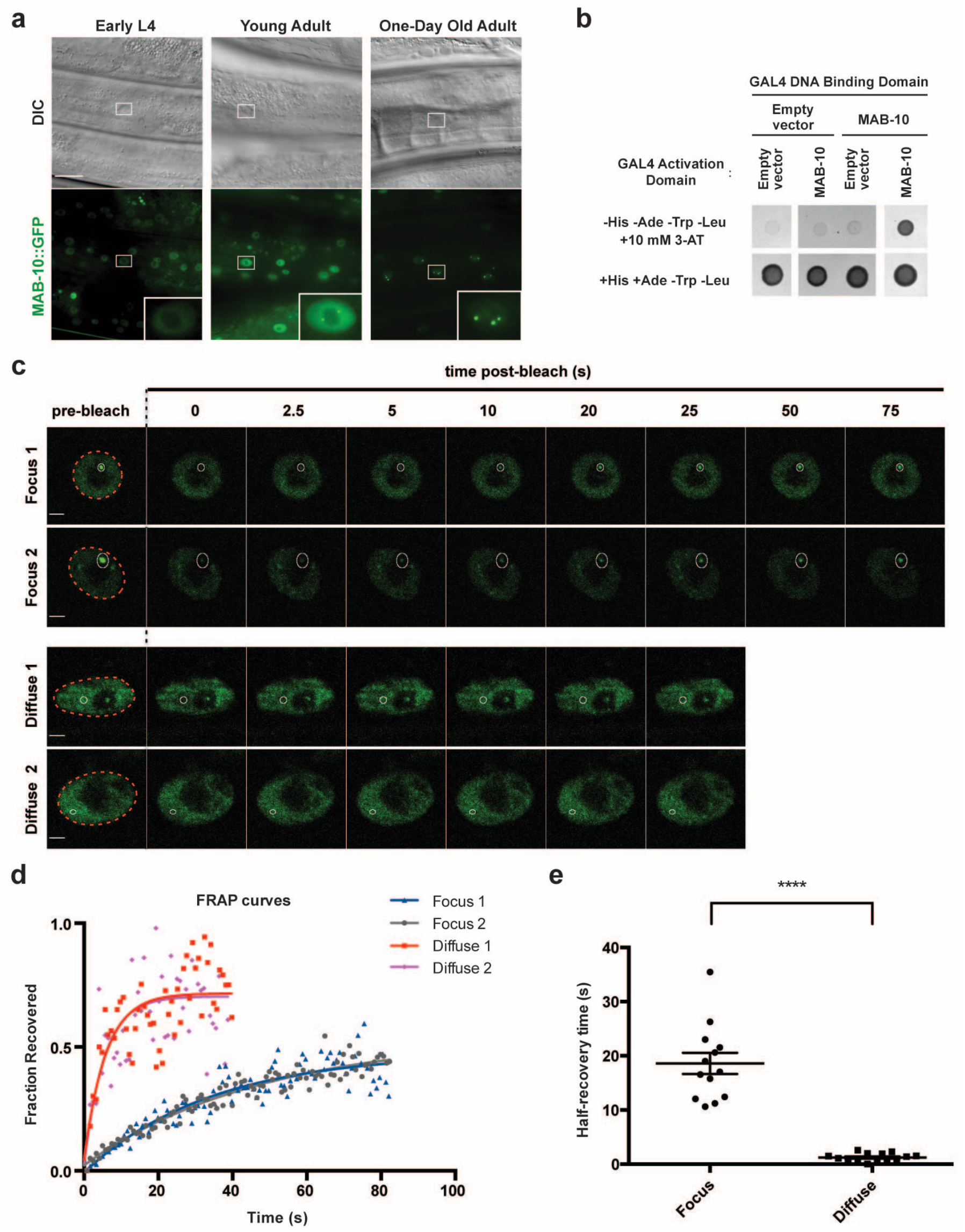
MAB-10 protein forms dynamic nuclear foci. **a**, Differential interference contrast (DIC) and confocal GFP fluorescence images of hypodermal cells in early L4, young adult, and one-day old adult wild-type animals expressing the translational reporter *n5909 [mab-10::gfp]*, which tags the endogenous *mab-10* gene locus with *gfp*. MAB-10::GFP expression in hypodermal cell nuclei begins in early L4 animals, increases and then starts forming nuclear foci in young adults; in one-day old adults, the majority of the MAB-10::GFP signal is condensed in nuclear foci. Scale bar, 20 μm. **b,** Yeast two-hybrid spot assay showing the self-interaction of MAB-10 protein (upper right spot) in quadruple drop-out plates (-His/-Ade/-Trp/-Leu) containing the competitive inhibitor of histidine synthesis 3-AT. **c,** Representative confocal fluorescent images of individual frames from *in vivo* FRAP time-lapse studies of two MAB-10 nuclear foci (“Focus”) and two diffuse MAB-10::GFP signals (“Diffuse”) in hypodermal nuclei of one-day old wild-type adults expressing the reporter *n5909 [mab-10::gfp].* Dotted red circle, nuclear circumference; white circle, focal area of laser bleaching and region of measurement; scale bars, 2 μm. **d,** FRAP curves quantifying observations from the experiments shown in (**c**) (two MAB-10 nuclear foci (“Focus”) and two diffuse MAB-10::GFP signals (“Diffuse”) in hypodermal nuclei). Fluorescence intensity measurements in the area of laser bleaching, normalized by considering the pre-bleach fluorescence intensity to be 1 and post-bleach intensity to be 0, are shown. t = 0, time measuring pre-bleaching intensity. Recovery of MAB-10::GFP in foci is significantly slower than in diffuse signal. **e,** Half-maximal recovery times from FRAP data for 13 experiments examining MAB-10::GFP in foci (“Focus”) and 14 experiments examining MAB-10::GFP diffuse signal (“Diffuse”). Error bars, SEM. ****p < 0.0001 (t test).

Protein self-interaction has been reported to be a general feature of proteins found in nuclear bodies^28, 29^. Using a yeast two-hybrid spot assay, we showed that MAB-10 proteins can self-interact (**Fig. 1b**), suggesting the role of MAB-10 self-association in its localization to nuclear foci. To examine the dynamic properties of MAB-10 nuclear foci, we performed fluorescence recovery after photobleaching (FRAP) studies and time-lapse imaging of MAB-10::GFP foci *in vivo* using one-day old adults (**Fig. 1c**). We found that MAB-10::GFP foci recovered fluorescence at a slower rate after photobleaching compared to that of diffuse MAB-10::GFP in the hypodermal nuclei in which the fluorescence was distributed throughout the nucleus (**Fig. 1c, d**). The half-maximal fluorescence recovery time, a measure of the MAB-10 protein exchange rate between the photobleached spot and the surrounding environment, was substantially longer for foci (*t*_1/2_ of ∼18.6 seconds) than for the diffuse MAB-10::GFP signal in the hypodermal nuclei (*t*_1/2_ of ∼1.2 seconds) (**Fig. 1e**). The reduced mobility of MAB-10 proteins in nuclear foci as revealed by our FRAP data suggests that diffusion barriers affect kinetic properties of the molecules within these structures.

### DDX-23 is a regulator of stem-cell fate in *C. elegans* hypodermal seam cells

While *lin-29* mutants fail at all of the events that characterize the larval-to-adult transition, *mab-10* mutants fail at only a subset of these events^6^ (**Supp Fig. 1a**), suggesting that there are factors that act with or in parallel to MAB-10 in controlling terminal seam-cell differentiation and the onset of adulthood. To identify such factors, we performed a mutagenesis screen using a reporter for the adult-specific *col-19* collagen gene, a downstream target gene of the heterochronic pathway that is highly expressed in the adult hypodermis^9, 30^. In wild-type adults the *col-19::GFP* fluorescent reporter (*maIs105*) is up-regulated, whereas in *lin-29* mutant adults this reporter is not expressed. In *mab-10* mutant adults *maIs105* reporter expression is reduced relative to that in wild-type animals (**Supp Fig. 1b**). We screened *mab-10(lf)* mutants for second-site mutations that further reduced their low *col-19* expression level (**Fig. 2a**). From this screen, we isolated two mutations (*n5703* and *n5705*) that proved to be alleles of the gene *ddx-23* (**Fig. 2b**). Strikingly, these *ddx-23* mutations alone caused dramatic reduction of *col-19* GFP expression in adult hypodermal cells in a wild-type background (**Fig. 2c**). Expressing a wild-type copy of the *ddx-23* gene (using *ddx-23(+)* transgenes) in worms with *ddx-23* mutations restored *col-19* expression (**Fig. 2b, c**). We generated the *ddx-23* allele *n6026* by recreating the I383F missense mutation of *ddx-23(n5705)* mutants using CRISPR-Cas9 and showed that this allele phenocopied the abnormal seam cell phenotype (**Supp Fig. 2a-d**). These observations established that *ddx-23* loss-of-function reduces *col-19* expression levels, and that *ddx-23*, like *lin-29*, is required for *col-19* up-regulation in the adult.

**Figure 2.**
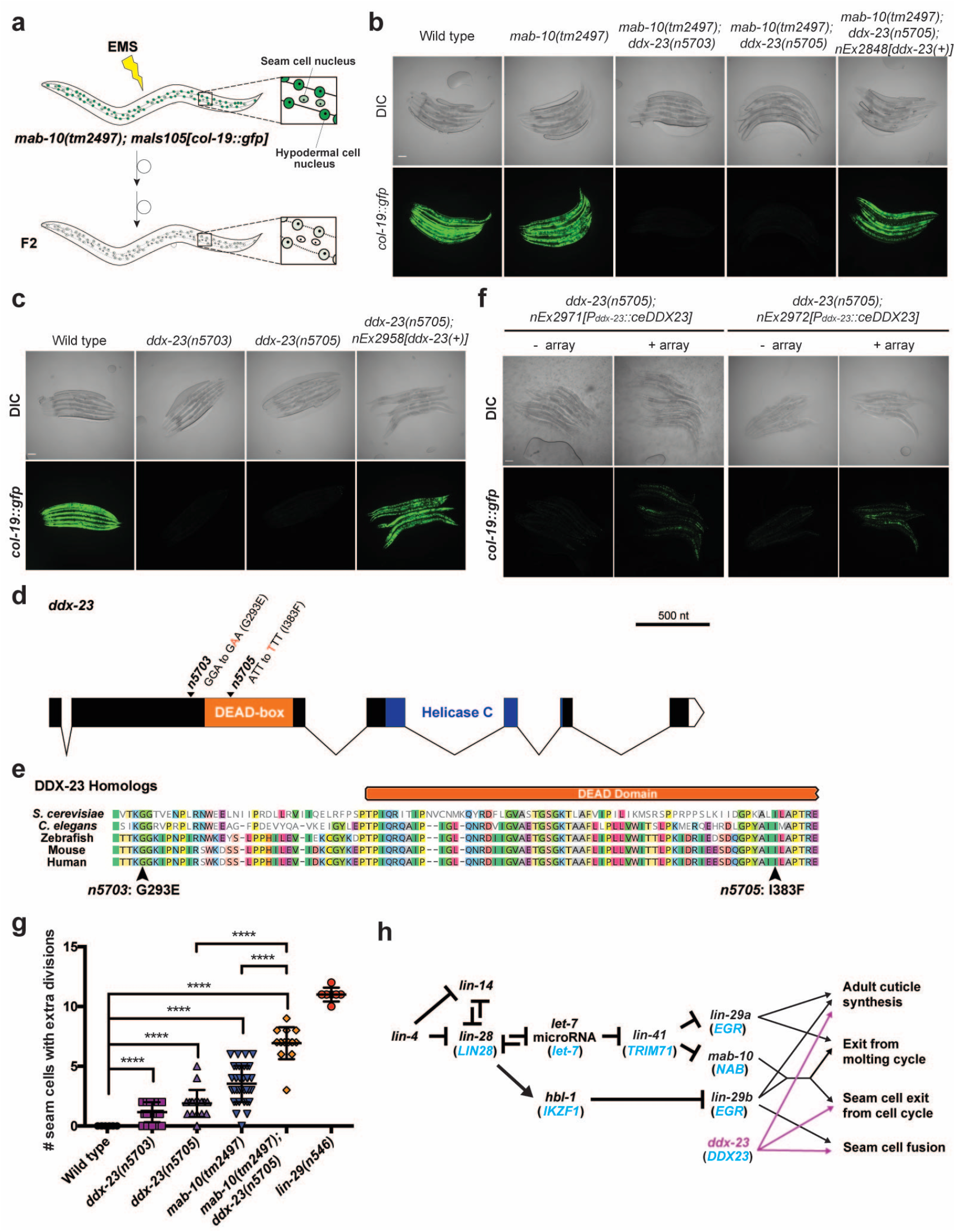
DDX-23 is evolutionarily conserved and functions in hypodermal stem cell biology. **a**, Schematic of the F2 non-clonal *mab-10* enhancer screen to identify factors that act with or in parallel to MAB-10. *mab-10(tm2497)* mutants were EMS-mutagenized, and F2 progeny were screened for second-site mutations that further reduced the low hypodermal *col-19* expression levels of *mab-10(tm2497)* mutants. **b,** The *mab-10* enhancer screen isolated two alleles (*n5703 and n5705)* of *ddx-23*. The *ddx-23(+)* transgene *nEx2848*, which contains wild-type *ddx-23*, rescued the low *maIs105* GFP expression. Representative DIC and GFP fluorescence micrographs are shown of *C. elegans* adults of the genotypes indicated and grown at 20 °C. Scale bar, 100 μm. **c,** Representative DIC and GFP fluorescence micrographs of wild-type and *ddx-23* single mutant adult animals grown at 20 °C. The *ddx-23(+)* transgene *nEx2958*, which contains wild-type *ddx-23*, rescued the low *maIs105* GFP expression. Scale bar, 100 μm. **d,** Schematic of *ddx-23* gene structure. The *ddx-23(n5703)* and *ddx-23(n5705)* mutations isolated from the *mab-10* enhancer screen are indicated. **e,** Sequence alignments of DDX-23 homologs from *S. cerevisiae* (Prp28), zebrafish (Ddx23), *Mus musculus* (DDX23), and *Homo sapiens* (DDX23). Only the region surrounding amino acid residues G293 and I383, which are altered by the *ddx-23* mutations *n5703* and *n5705,* respectively, is shown. Arrows indicate the completely conserved G293 and I383 residues. **f,** Representative micrographs of *maIs105[col-19::gfp]* reporter GFP expression in one-day old adults of the indicated genotypes. Overexpression of human *DDX23* codon-optimized for expression in *C. elegans (ceDDX23)* partially rescued the abnormal *maIs105[col- 19::gfp]* GFP expression seen in *mab-10(tm2497); ddx-23(n5705)* animals. Scale bar, 100 μm. **g,** Number of seam cells that underwent extra seam cell division, scored in one-day old adults. Error bars, SD. ****p < 0.0001 (t test). **h,** Proposed role of the *ddx-23* gene in the heterochronic pathway. DDX-23 acts with MAB-10 to control seam cell exit from the cell cycle but acts independently of MAB-10 to control seam cell fusion in adults. The human homologs of the *C. elegans* heterochronic genes are indicated in blue.

The *ddx-23* gene encodes an evolutionarily conserved DEAD-box helicase protein (**Supp Fig. 1c**) that is generally involved in RNA biogenesis. In *C. elegans*, DDX-23 has been shown to regulate tissue differentiation^31^ and support microRNA biogenesis^32^. The two mutations we isolated are missense mutations (G293E and I383F) of amino acid residues that are highly conserved among DDX-23 homologs (**Fig. 2d, e**).

To ascertain if human DDX23 functions similarly to *C. elegans* DDX-23, we asked if expression of a transgene encoding the human DDX23 and codon-optimized for *C. elegans* expression (*ceDDX23)* could rescue the low *col-19* expression phenotype of *ddx-23* mutants. *ceDDX23* transgenic lines partially rescued this defect in *ddx-23* mutants (**Fig. 2f**). This result indicates that the *C. elegans ddx-23* and human *DDX23* genes likely function similarly and act in similar molecular genetic pathways, highlighting the evolutionarily conserved nature of *ddx-23* in stem-cell fate determination.

Because *col-19* encodes an adult-specific collagen and low *col-19* expression in *ddx-23* mutants (**Fig. 2c**) suggests a defect in the larval-to-adult transition, we asked if *ddx-23* mutants are defective in other aspects of the onset of adulthood in the wild type. We first examined the formation of adult alae, distinct lateral longitudinal cuticular structures, using electron microscopy. *ddx-23* mutants had partially disrupted alae (**Supp Fig. 2e**), consistent with previous findings that the disruption of COL-19 leads to defects in adult-specific alae structure^33^.

Furthermore, like *mab-10* mutants, *ddx-23* mutants had seam cells that underwent extra divisions, and a *ddx-23* mutation enhanced the supernumerary-seam-cell phenotype of *mab-10* mutants (**Fig. 2g**). Additionally, in *ddx-23* mutant adults some seam cells failed to undergo the terminal fusion characteristic of the larval-to-adult transition of wild-type animals (**Supp Fig. 2f**). Like *ddx-23* single mutants, *mab-10 ddx-23* double mutants also showed a defect in seam cell fusion. *mab-10* does not function in this terminal fusion process^6^. We conclude that *ddx-23* plays multiple roles in the heterochronic pathway and the larval-to-adult transition, working with *mab-10* to control seam cell exit from the cell cycle while also functioning independently of *mab-10* in regulating seam cell fusion and adult alae formation (**Fig. 2h**, **Supp Fig. 1a**).

### DDX-23 controls MAB-10 nuclear foci formation

To examine how DDX-23 and MAB-10 function together to control seam cell exit from the cell cycle, we tested whether the *C. elegans* MAB-10 protein can physically interact with DDX-23 in yeast two-hybrid spot assays. We observed interactions between MAB-10 and DDX-23 (**Fig. 3a**). To determine if DDX-23 interacts with MAB-10 *in vivo* in *C. elegans*, we used a split GFP (spGFP) system^34, 35^ (**Fig. 3b**). We tagged DDX-23 with the N-terminal fragment of GFP (spGFPN) and MAB-10 with the C-terminal portion of GFP (spGFPC), and drove the expression of both by the seam cell promoter *P_ceh-16_*. GFP fluorescent signal should be detected only when the proteins containing the two spGFP fragments physically interact with each other. We found that DDX-23 interacted with MAB-10 *in vivo* in living *C. elegans* (**Fig. 3c**); no GFP signal was observed when an empty spGFPC control tag was used. This spGFP experiment also allowed us to identify the subcellular locations where DDX-23 and MAB-10 proteins interact.

**Figure 3.**
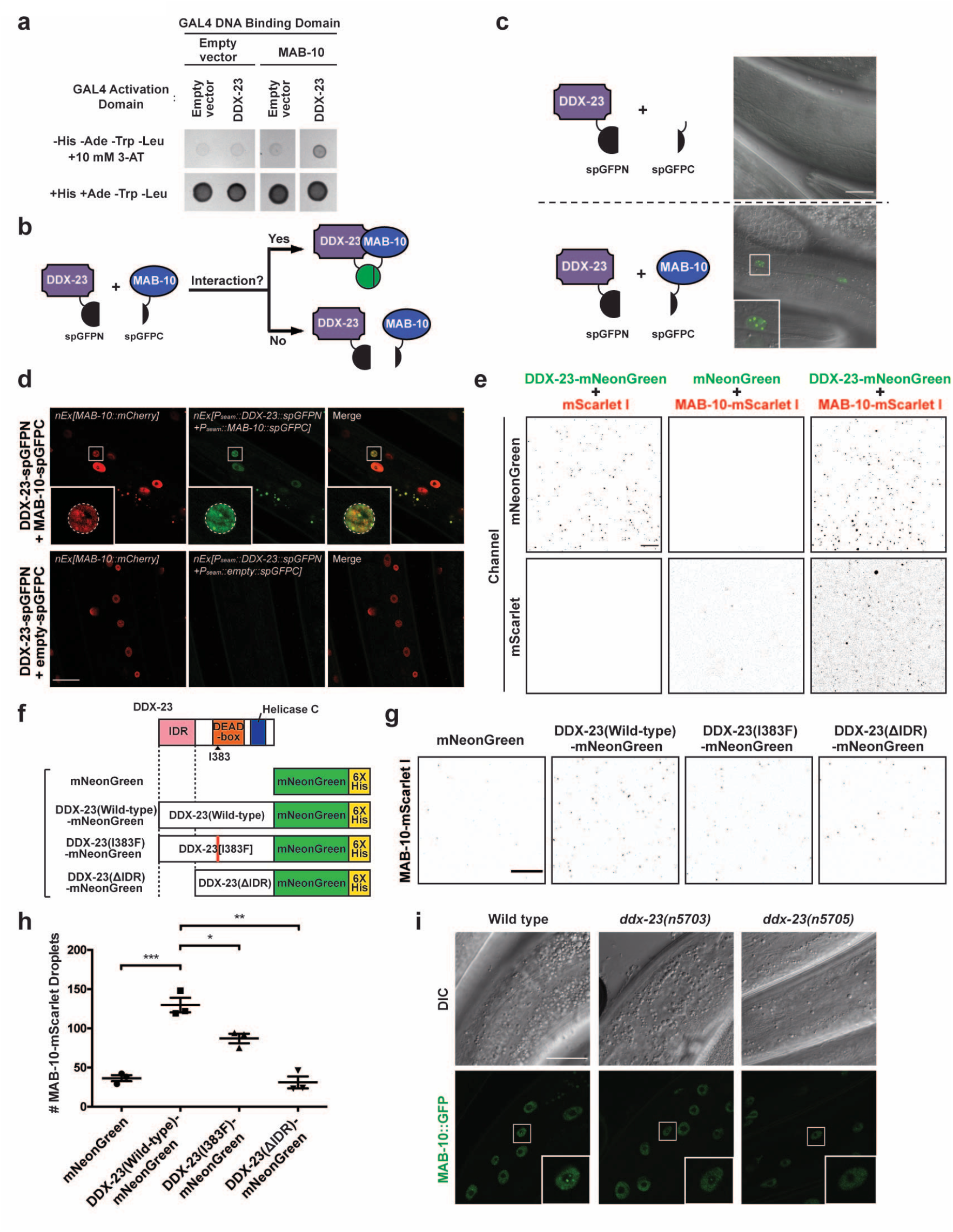
DDX-23 enhances the partitioning of MAB-10 proteins into nuclear foci. **a**, Yeast two-hybrid spot assay showing the interaction of DDX-23 protein fused to the GAL4 activation domain (upper right spot) but not of a control with the GAL4 activation domain-only (“Empty vector”) protein with MAB-10 protein fused to the GAL4 DNA-binding domain in quadruple drop-out plates (-His/-Ade/-Trp/-Leu) containing the competitive inhibitor of histidine synthesis 3-AT. **b,** Schematic representation of the split GFP (spGFP) approach for assessing protein-protein interactions *in vivo*. Animals expressed an N-terminal GFP fragment (spGFPN) fused to DDX-23 in combination with a C-terminal GFP fragment (spGFPC) fused to MAB-10. A physical interaction between the DDX-23::spGFPN and MAB-10::spGFPC proteins leads to the emission of green fluorescence. **c,** Representative confocal images (merged DIC and GFP channels) of spGFP experiments using DDX-23::spGFPN and MAB-10::spGFPC (or spGFPC by itself as a control). GFP fragments were expressed under the control of the seam cell promoter *P_ceh-16_*. Physical interaction, monitored by GFP signal, was detected only when DDX-23::spGFPN was combined with MAB-10::spGFPC. Scale bar, 20 μm. **d,** Confocal fluorescence images of animals expressing a MAB-10 translational reporter (*P_mab-10_::mab-10::mCherry::mab-10 3’UTR)*, DDX-23::spGFPN and MAB-10::spGFPC (or empty spGFPC as a control). Inset: seam cell. DDX-23 and MAB-10 interact at nuclear foci. Dotted white circle, nuclear circumference; scale bar, 20 μm. **e,** Representative images of *in vitro* assays testing heterotypic interactions of purified DDX-23-mNeonGreen and MAB-10-mScarlet I (right panel pairs). DDX-23-mNeonGreen can self-interact in the presence of mScarlet I alone (left panel pairs). MAB-10-mScarlet I can self-interact in the presence of mNeonGreen alone (middle panel pairs) but interaction is enhanced in the presence of DDX-23-mNeonGreen (right panel pairs). Micrographs were color-inverted for better visualization. Assays were performed in the presence of 220 mM NaCl using 500 nM of each purified protein (no PEG-8000). Scale bar, 100 μm. **f,** Schematic view of control mNeonGreen protein alone, wild-type DDX-23-mNeonGreen, mutant DDX-23(I383F)-mNeonGreen, and a DDX-23 protein lacking its IDR (DDX-23(ΔIDR)-mNeonGreen) used for recombinant protein production. **g,** Representative images of *in vitro* interaction assays of control mNeonGreen protein alone, wild-type DDX-23-mNeonGreen, mutant DDX-23(I383F)-mNeonGreen, and DDX-23(ΔIDR)-mNeonGreen, each with purified MAB-10-mScarlet I. Micrographs were color-inverted for better visualization. Assays were performed in the presence of 200 mM NaCl and 10 % PEG-8000 using 10 nM of each protein. Scale bar, 50 μm. **h,** Quantification of the number of MAB-10 spots when mixed in heterotypic assays with an mNeonGreen alone control, wild-type DDX-23, mutant DDX-23[I383F] or DDX-23(ΔIDR). Error bars, SEM. ***p<0.001, **p < 0.01 and *p < 0.05 (t test). **i,** DIC and confocal fluorescent images of hypodermal cells in one-day old wild-type, *ddx-23(n5703)* and *ddx-23(n5705)* adults expressing the reporter *n5909 [mab-10::gfp]*. Scale bar, 20 μm.

We observed diffuse spGFP signal in the seam cell nuclei along with distinct nuclear foci with concentrated spGFP fluorescence within these nuclei (**Fig. 3c, inset**). We then asked if these nuclear foci at which DDX-23 and MAB-10 interact are the same sites at which MAB-10 condensates form. We generated a transgenic reporter strain that expresses an *mCherry* translational reporter for MAB-10 (*P_mab-10_::mab-10::mCherry*) as well as the spGFP constructs that we used to observe the interaction between DDX-23 and MAB-10 (DDX-23-spGFPN and MAB-10-spGFPC). We showed by co-localization of the mCherry and GFP signals that the MAB-10 nuclear foci are indeed the sites of interaction between

DDX-23 and MAB-10 (**Fig. 3d**). We note that the addition of the *mCherry* translational reporter for MAB-10 (*P_mab-10_::mab-10::mCherry*) to the spGFP experiment reduced the brightness and distinct nature of the DDX-23-MAB-10 spGFP foci; we suggest that the overexpressed MAB-10::mCherry from the extrachromosomal array competed with MAB-10::spGFPC for the binding to DDX-23::spGFPN.

Both DDX-23 and MAB-10 proteins contain large intrinsically disordered regions (IDRs) (**Supp Fig. 3a**). Interactions between IDRs can act as a driving force for protein condensation into sub-nuclear bodies^36^. We first asked if DDX-23 and MAB-10 can form homotypic interactions *in vitro* and found that both purified recombinant DDX-23-mNeonGreen fusion protein (**Supp Fig. 3b, c, d, e**) and MAB-10 - mScarlet I fusion protein (**Supp Fig. 3f, g, h**) showed homotypic interactions.

Since our *in vivo* data showed that DDX-23 and MAB-10 proteins interact at nuclear foci (**Fig. 3c, d**), we asked whether the DDX-23 protein influences the incorporation of MAB-10 protein into these nuclear structures. We assessed the *in vitro* formation of MAB-10 - mScarlet I protein condensates in the presence or absence of a DDX-23 - mNeonGreen fusion protein. The MAB-10 protein condensed more efficiently (∼3.5 fold, p < 0.001) in the presence of DDX-23 (**Fig. 3e, f, g, h**), indicating that DDX-23 can facilitate the condensation of MAB-10 protein. As shown above, the DDX-23 missense mutation I383F causes a defect in seam cell fate (**Fig. 2, Supp Fig. 2**). We asked if this mutation can influence MAB-10 foci formation by testing the ability of MAB-10 to be incorporated into MAB-10 structures in a heterotypic assay with DDX-23[I383F] protein. The efficiency of MAB-10 incorporation into MAB-10 structures *in vitro* was reduced by ∼33 % (p < 0.05) in the presence of mutant DDX-23[I383F] protein as compared to wild-type protein (**Fig. 3f, g, h**). We next asked how the IDR of DDX-23 might influence MAB-10 droplet formation. Specifically, using heterotypic assays of MAB-10 and DDX-23 proteins, we observed a 76% reduction (p < 0.01) in the number of MAB-10 structures formed in assays involving a deletion of DDX-23 lacking its IDR (DDX-23(ΔIDR), **Fig. 3f**) compared to the number formed in assays performed with wild-type DDX-23 (**Fig. 3g, h**). These results suggest that in addition to the DEAD-box domain the low-complexity IDR of DDX- 23 is also involved in driving the condensation of MAB-10.

We examined the ability of DDX-23 to facilitate the formation of MAB-10 foci *in vivo* by comparing MAB-10::GFP foci in wild-type and *ddx-23* mutants. We found that for the *n5703* and *n5705* alleles of *ddx-23* the number of animals with MAB-10::GFP foci in adult hypodermal nuclei was reduced by 60% (p = 0.0002) and 76% (p < 0.0001), respectively (*n* = 30 for each genotype)(**Fig. 3h**). This result indicates that wild-type DDX-23 protein functions *in vivo* to facilitate MAB-10 condensate formation in the nuclei of hypodermal and seam cells in adult animals.

### MAB-10 and DDX-23 transcriptionally repress larval genes, including *hedgehog*-related genes

We examined the roles of LIN-29, MAB-10 and DDX-23 in regulating gene expression. Since MAB-10 functions as a transcriptional cofactor for LIN-29 to regulate a subset of LIN-29 target genes and our data showed that the adult-specific MAB-10 nuclear foci formation was in part facilitated by DDX-23, we performed RNA sequencing (RNA-Seq) studies of *lin-29*, *mab-10*, and *ddx-23* mutant adults and compared their gene expression profiles to that of wild-type adults. We found that the majority of genes that were misregulated in all three mutants were up-regulated (**Fig. 4a**), showing a major role for LIN-29, MAB-10, and DDX-23 proteins in transcriptional repression. Importantly, the set of genes co-repressed by LIN-29, MAB-10, and DDX-23 was enriched for genes known to be involved in the larval-to-adult transition. For example, many larval-specific collagen genes are normally expressed highly in L4 stage animals and down-regulated in adults; these genes were up-regulated in *lin-29*, *mab-10*, and *ddx-23* mutant adults compared to wild-type adults (**Supp Fig. 4**). This observation indicates that LIN-29, MAB-10, and DDX-23 proteins are critical in establishing an adult gene expression profile by transcriptionally repressing larval-related genes.

**Figure 4.**
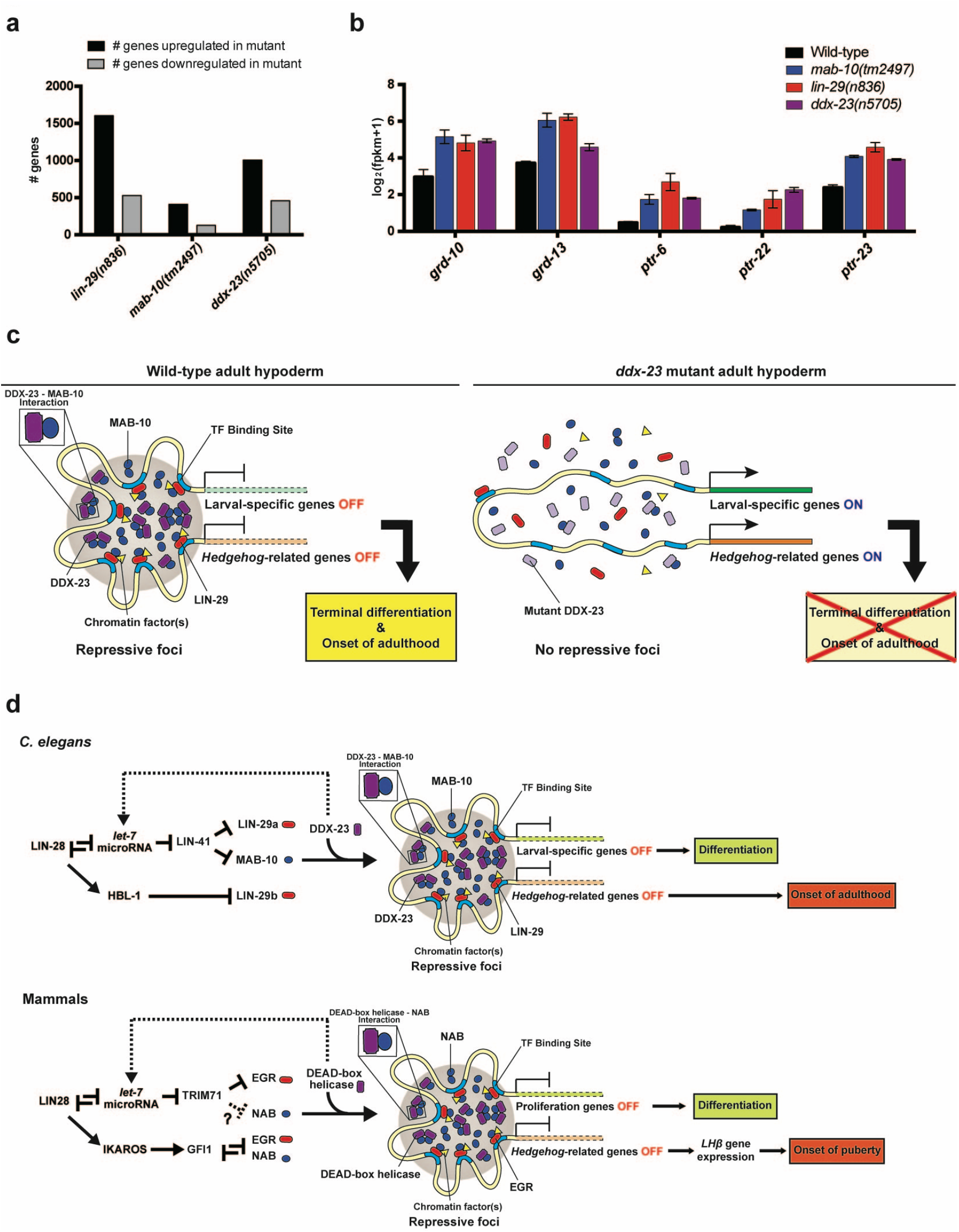
DDX-23-driven formation of MAB-10 (NAB) nuclear foci might control terminal differentiation and the onset of adulthood by transcriptionally repressing LIN-29 (EGR) target *hedgehog* and larval-specific genes. **a**, RNA-Seq analysis of *lin-29*, *mab-10*, and *ddx-23* mutant adults. Number of genes up-regulated (black) and down-regulated (gray) relative to wild-type adults are shown for the indicated genotypes. **b,** Normalized gene expression abundance, calculated as log2(fragments per kilobase million (fpkm) +1), of several *hedgehog*-related (*grd)* genes and their putative receptor (*ptr*) genes that are up-regulated in *lin-29*, *mab-10*, and *ddx-23* mutant adults, as identified by RNA-Seq analysis. Error bars, SEM. p < 0.01 for all comparisons between wild-type vs. mutant. **c,** A proposed model for how DDX-23-driven formation of MAB-10 nuclear foci controls LIN-29-regulated target gene repression in the adult hypoderm. DDX-23 binds to and enhances the partitioning of MAB-10 proteins into foci, causing repression of LIN-29-regulated target genes -- including larval-specific and *Hedgehog*-related genes -- and thereby driving terminal cell differentiation and the onset of adulthood (left panel). Loss-of- function mutations in *ddx-23* reduce MAB-10 foci and up-regulate larval-specific and *Hedgehog*-related genes, resulting in a failure of the seam cells to terminally differentiate and of transition to adulthood (right panel). **d,** Models for how the heterochronic pathway regulates cell differentiation and the onset of puberty in *C. elegans* (upper panel) and mammals (lower panel). For example, LIN28 signaling in mammals (LIN-28 signaling in *C.* elegans) coordinates the expression of EGR (*C. elegans* LIN-29) and NAB (*C. elegans* MAB-10) proteins, both *let-7* microRNA-dependently via TRIM71 (mammalian homolog of the *C. elegans* LIN-41) and *let-*7-independently via IKAROS (mammalian homolog of the *C. elegans* HBL-1) and GFI1. The formation of nuclear foci that contain NAB (MAB-10) proteins is facilitated by a DEAD-box helicase protein (DDX-23) and function to transcriptionally repress EGR (LIN-29) target genes including proliferation-related genes and *hedgehog*-related genes. The repression of proliferation-related genes leads to cellular differentiation, while *Hedgehog* signaling leads to luteinizing hormone LHβ expression and the onset of puberty. DDX-23 was previously described to play a role in the primary microRNA processing of *let-7* microRNA^32^ (dotted lines), possibly providing a feedback regulatory loop in this pathway.

Strikingly, we also found several *hedgehog*-related genes and genes that encode putative *hedgehog* receptors to be up-regulated in *lin-29, mab-10*and *ddx-23* mutant strains (**Fig. 4b**). These genes are the evolutionary homologs of *Hedgehog (Hh)* and its receptor *Patched* (*Ptc*), which control various developmental processes in species ranging from *Drosophila* to humans^37^ and have been implicated in the regulation of mammalian stem cell maintenance and proliferation^38, 39^. *Hedgehog*- and *Patched*-related genes are known to be involved in *C. elegans* molting^40^ but have not been previously linked to the heterochronic pathway, which includes *lin-29, mab-10* and now *ddx-23*.

## Discussion

Recent studies of transcriptional condensates have revealed new possible mechanisms of transcriptional control. There is growing evidence that transcriptional condensates are involved in both the activation^1–5, 41–43^ and the repression^44–47^ of gene expression. However, little is known about how the formation of such condensates is controlled. In this study, we report that *in vivo* the *C. elegans* DEAD-box helicase protein DDX-23 regulates stem-cell fate at least in part by binding to and enhancing the partitioning of MAB-10 proteins into condensates, which in turn repress LIN-29-regulated target genes -- including known larval-specific genes and *Hedgehog*-related genes -- and thereby drive terminal cell differentiation and the onset of adulthood (**Fig. 4c**). Loss-of-function mutations in *ddx-23* reduce MAB-10 repressive condensates and up-regulate larval-specific genes and *Hedgehog*-related genes, resulting in a failure of the seam cells to terminally differentiate and transition to adulthood (**Fig. 4c**).

Previous studies of *C. elegans*, mice and human cell lines have shown that suppression of LIN-28 levels promotes the expression of LIN-29 (EGR) and MAB-10 (NAB) proteins, both *let-7* microRNA-dependently via LIN-41 (mammalian homolog TRIM71)^21–23^ and *let-*7-independently via HBL-1 (mammalian homolog IKAROS)^23, 48–50^ and GFI1^51^ (**Fig. 4d**). The *ddx-23* gene was previously found to play a role in the primary microRNA processing of *let-7* microRNA^32^. Here we describe a novel gene regulatory mechanism that regulates the activities of MAB-10 (NAB) and LIN-29 (EGR): DDX-23 drives the condensation of MAB-10 (NAB) protein into nuclear focal structures to spatiotemporally organize the nucleus and transcriptionally repress LIN-29 (EGR) target larval genes, including *hedgehog*-related genes. We propose that this process is evolutionarily conserved and that a comparable molecular genetic mechanism -- in which the partitioning of NAB proteins by a DEAD-box helicase drives repression of proliferation-related and *Hedgehog*-related genes by EGR -- controls mammalian stem cell development and the onset of puberty (**Fig. 4d**). In mammals, the DEAD-box helicase DDX5 protein^52^ and EGR1 protein^16^ act as a barrier to somatic cell reprogramming^52^, and the DEAD-box protein Ddx20/DP103 represses transcriptional activation by Egr2^56^. We propose that DEAD-box proteins function to control EGR transcriptional activity by facilitating the condensation of its cofactor NAB, thereby coordinating molecular genetic programs that control cell differentiation.

The highly conserved *LIN28/let-7* axis has previously been found to regulate *Hedgehog* signaling in multiple contexts, *e.g*., during retina regeneration in zebrafish^54^ and in a mouse model for embryonal tumors with multilayered rosettes (ETMRs), a tumor type characterized by LIN-28A overexpression^55^. *LIN28* is implicated in regulating the onset of puberty in mice and humans^18, 56–60^, while *Hedgehog* signaling controls cell proliferation and cell-type determination in the pituitary gland of mice by regulating the luteinizing hormone LHβ expression, a key hormone involved in the onset of puberty^61–64^. Our findings might provide a missing link between the LIN28/*let-7* axis and *Hedgehog* signaling: LIN28 signaling promotes the expression of EGR and NAB proteins, which in turn leads to the repression of *Hedgehog* genes by EGR. This repression is achieved by the condensation of NAB proteins to form a repressive condensate, a process driven in part by a DEAD-box helicase protein. The repression of *Hedgehog-related* genes subsequently induces the expression of the luteinizing hormone LHβ and leads to cell differentiation and the onset of puberty (**Fig. 4d**).

Our findings have important implications for the understanding of molecular mechanisms of carcinogenesis. The overexpression of human *LIN28*inhibits *let-7* microRNA gene activity and thereby promotes cancerous growth^13, 65^, possibly by augmenting downstream *Shh* Hedgehog signaling^55^. We speculate that this regulation is driven by the down-regulation of EGR, which acts as a tumor suppressor and is commonly mutated or suppressed in multiple human cancers^66, 67^ and that this down-regulation leads to the up-regulation of EGR target genes (including *Hedgehog* genes) that are normally silenced to regulate cell differentiation and the onset of adulthood. In healthy cells, DEAD-box helicase proteins ensure repression of EGR target genes essential to cell proliferation by facilitating the formation of MAB-10 (NAB) repressive transcriptional condensates; failure to form such condensates could lead to abnormal cell proliferation. Indeed, *DDX23* mutations and deletions have been detected in several human cancers^68^, suggesting a tumor suppressor role for this gene. Based on our findings, we propose as a mechanism that these *DDX23* genetic defects result in a failure to form DDX23-NAB transcriptional condensates and the consequent up-regulation of EGR-repressed targets -- which include genes involved in *Hedgehog* signaling – leading to the abnormal cell proliferation that drives cancerous growth.

---

**Supplementary Figure 1.**
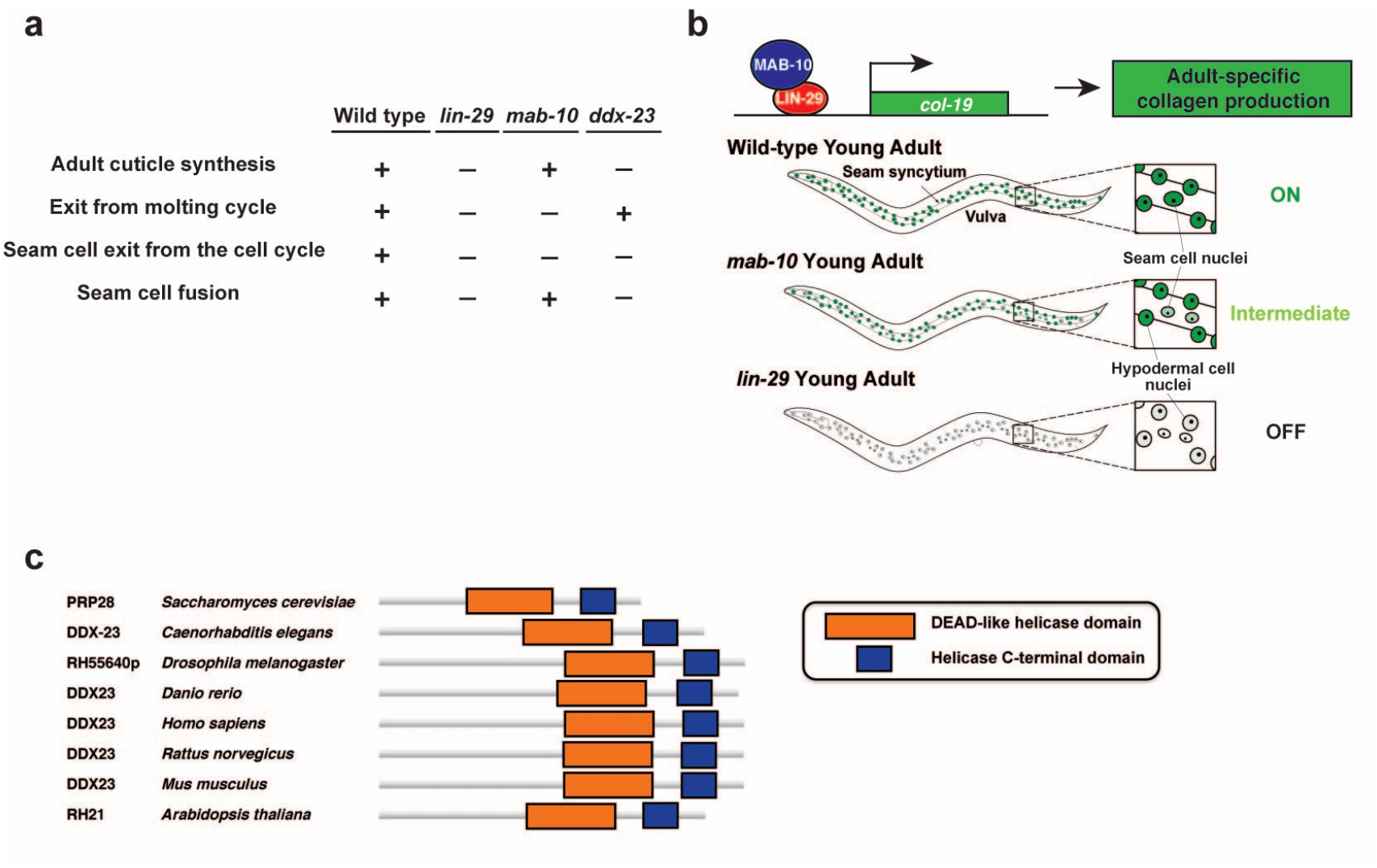
*lin-29* and *mab-10* mutant phenotypes and molecular identification of *ddx-23*. **a**, Presence(+) or absence(-) of the four key characteristic events that normally occur during the larval-to-adult transition for the indicated genotypes. **b,** Rationale for the F2 non-clonal screen for factors that act with or in parallel to MAB-10. The *col-19::gfp (maIs105)* transgene was used as a reporter for the heterochronic pathway. In wild-type animals *maIs105* is up-regulated in adults, whereas in *lin-29* mutant adult animals this reporter is not expressed. In *mab-10* mutant adult animals there is a reduced level of *maIs105* expression. **c,** Homologs of the *C. elegans* DDX-23 protein contain the DEAD-like helicase domain (orange) and the helicase C-terminal domain (blue).

**Supplementary Figure 2.**
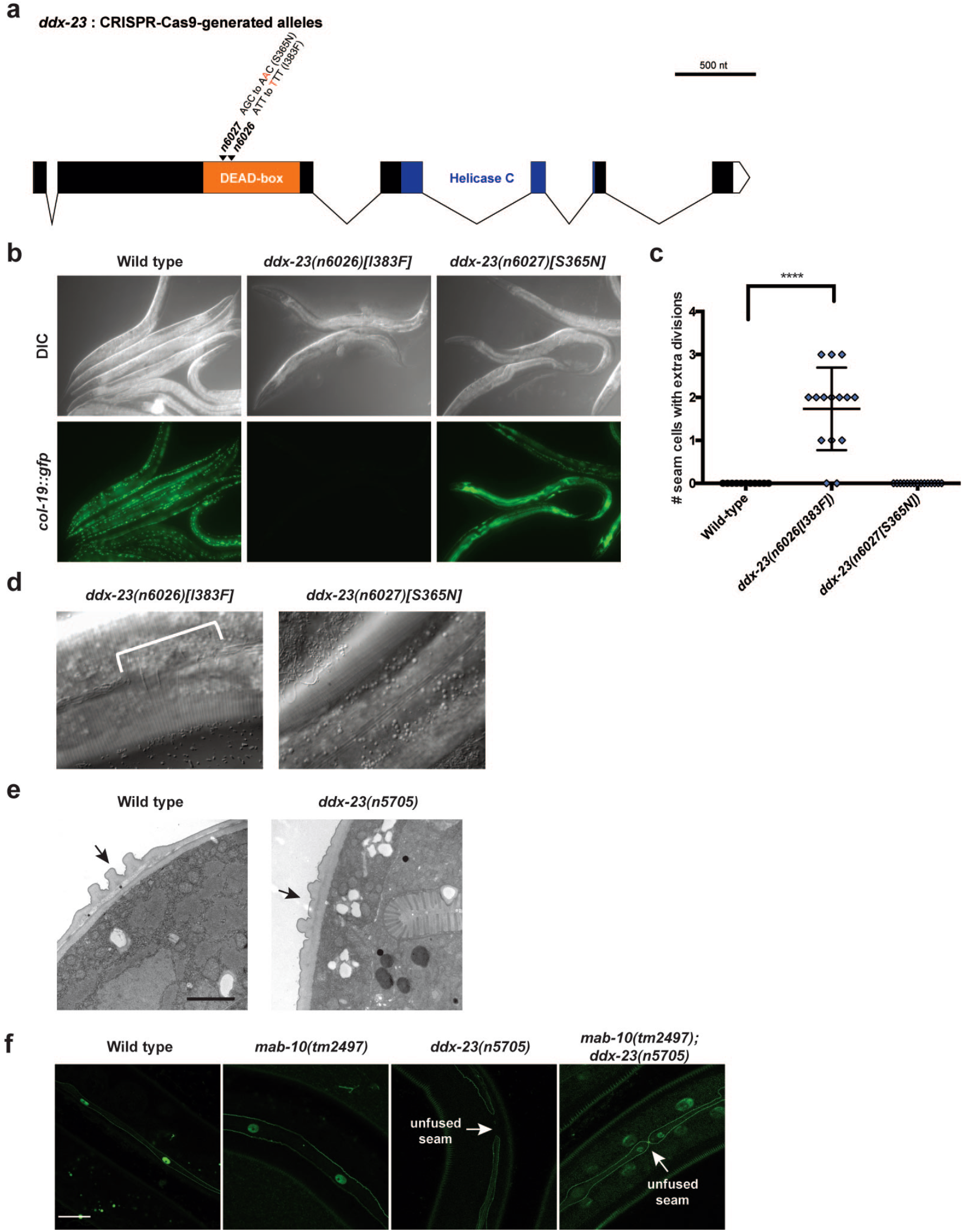
CRISPR-Cas9-generated *ddx-23(n6026)* allele encoding DDX-23[I383F] phenocopies the *ddx-23(n5705)* abnormal phenotype. **a**, Schematic of *ddx-23* gene structure. *ddx-23(n6026)[I383F]* and *ddx-23(n6027)[S365N]* alleles were individually recreated using the CRISPR-Cas9 genome-editing technology. Both of these DDX-23 missense mutations (S365N, I383F) were present in the strain isolated from the *mab-10*enhancer screen. For simplicity, we designated the allele *ddx-23(n5705)* from the *mab-10* enhancer screen as the causal mutation for the abnormal phenotypes and concluded that the DDX-23[I383F] mutation is responsible for the phenotypic effects based on the observation that *ddx-23(n6026)[I383F]* but not *ddx-23(n6027)[S365N]* phenocopied the *ddx-23(n5705)* seam cell abnormalities (Figure 2c, g and Extended Data Figure 2b-d). **b,** Representative micrographs of *maIs105[col-19::gfp]* reporter expression in one-day old adults of the indicated genotypes. *ddx-23(n6026)[I383F]* but not *ddx-23(n6027)[S365N]* mutants showed low *col-19* transgene expression. **c,** Number of seam cells that undergo extra seam cell division, scored in one-day old adults. *ddx-23(n6026)[I383F]* but not *ddx-23(n6027)[S365N]* mutants had seam cells that underwent extra divisions. Error bars, SD. ****p < 0.0001 (t test). **d,** Representative micrographs of the lateral alae, longitudinal cuticular ridges generated by the seam cells, of *ddx-23(n6026)[I383F]* and *ddx-23(n6027)[S365N]* one-day old adults. *ddx-23(n6026)[I383F]* but not *ddx-23(n6027)[S365N]* mutants had defective alae formation. **e,** Electron micrographs of cross-sections of wild-type and *ddx-23(n5705)* 24 hour adult males. Arrows indicate adult-specific lateral alae. Scale bars, 2 μm. **f,** Confocal fluorescence images of seam cell fusion events observed in one-day old adults of the indicated genotypes containing the reporter *wIs78[ajm-1::gfp + scm::gfp]*, which marks the seam cell nuclei and adherens junctions. Scale bar, 20 μm.

**Supplementary Figure 3.**
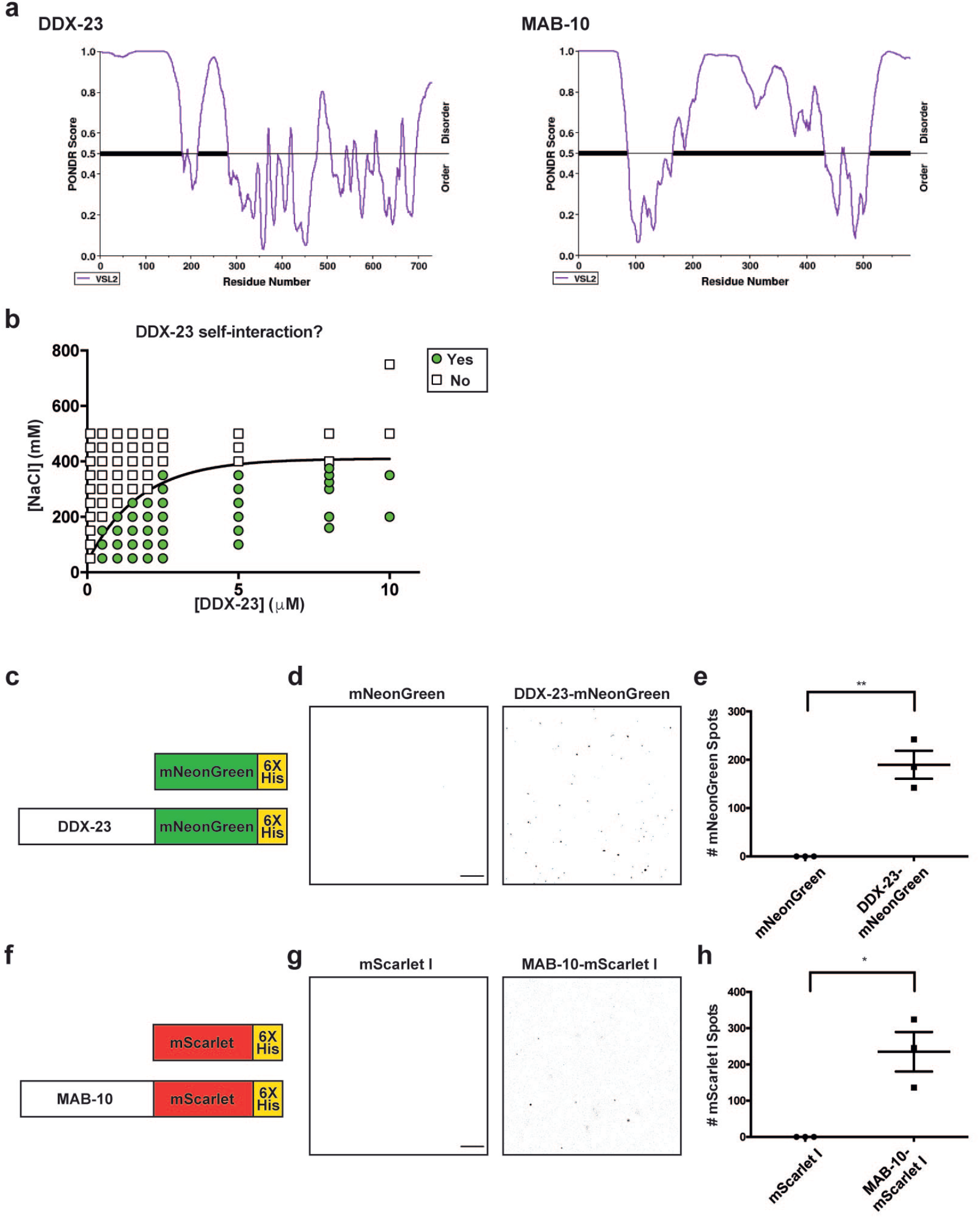
DDX-23 and MAB-10 self-interact. **a**, Graphs plotting intrinsic disorder of DDX-23 (left panel) and MAB-10 (right panel). PONDR (Predictor of Natural Disordered Regions) VSL2 scores^69^ (http://www.pondr.com/) are shown on the y axis, and amino acid positions are shown on the x axis. PONDR scores show predictions of ordered (PONDR score < 0.5) or disordered (PONDR score > 0.5) structure throughout the protein. **b,** Diagram plotting purified DDX-23 protein and NaCl concentrations, scoring for presence (green circles) or absence (white squares) of optically resolvable spots. Assays were performed in the absence of the molecular crowder PEG-8000^70^ to determine if the DDX-23 protein is capable of self-interacting under physiologically relevant concentrations. **c,** Schematic view of control mNeonGreen protein and DDX-23-mNeonGreen used for recombinant protein production. **d,** Representative images of *in vitro* assays testing homotypic interaction of purified DDX-23-mNeonGreen (right panel) and mNeonGreen alone as a control (left panel). Micrographs were color-inverted for better visualization. Assays were performed in the presence of 220 mM NaCl using 500 nM purified protein (no PEG-8000). Scale bar, 100 μm. **e,** Number of spots of control mNeonGreen alone and of DDX-23-mNeonGreen. Error bars, SEM. **p < 0.01 (t test). **f,** Schematic view of control mScarlet I protein and MAB-10-mScarlet I used for recombinant protein production. **g,** Representative images of *in vitro* assays testing homotypic interaction of purified MAB-10-mScarlet I (right panel) and mScarlet I alone as a control (left panel). Micrographs were color-inverted for better visualization. Assays were performed in the presence of 220 mM NaCl using 500 nM purified protein (no PEG-8000). Scale bar, 100 μm. **h,** Number of homotypic spots of control mScarlet I alone and of MAB-10-mScarlet I. Error bars, SEM. *p < 0.05 (t test).

**Supplementary Figure 4.**
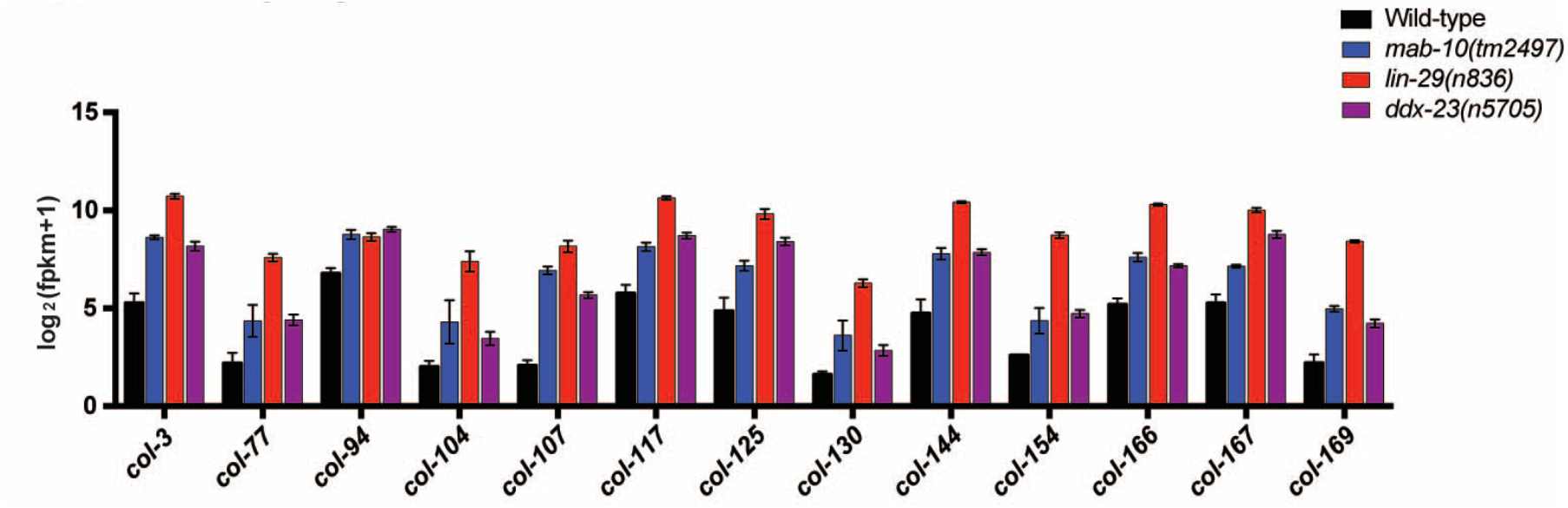
LIN-29, MAB-10, and DDX-23 proteins are critical in establishing an adult gene expression profile by transcriptionally repressing larval-related genes. Normalized gene expression abundance, calculated as log_2_(fragments per kilobase million (fpkm) +1), of multiple collagen genes that are up-regulated in *lin-29*, *mab-10*, and *ddx-23* mutant adult animals, as identified by RNA-Seq analysis. In wild-type animals, these collagen genes are expressed highly in L4 stage animals and decrease in expression in adults. Error bars, SEM. p < 0.01 for all comparisons between wild-type vs. mutant.

## Contact for Reagent and Resource Sharing

Further information and requests for resources and reagents should be directed to and will be fulfilled by the Lead Contact, H. Robert Horvitz.

## Materials and Methods

### *C. elegans* strains and transgenes

All *C. elegans* strains were cultured as described previously^71^. We used the N2 Bristol strain as the reference wild-type strain, and the polymorphic Hawaiian strain CB4856 for genetic mapping and SNP analysis^72^. All strains were maintained at 20 °C.

We used the following mutations and transgenes:

LGII: *mab-10(tm2497, n5909 [mab-10::gfp]), lin-29(n546, n836)*

LGIII: *ddx-23(n5703, n5705, n6026[I383F], n6027[S365N])*

LGIV: *wIs78 [ajm-1::gfp + scm::gfp]*

LGV: *maIs105[col-19::gfp]*

### Extrachromosomal arrays

*nEx2848[P_ddx-23_::ddx-23::ddx-23 3’UTR, P_ttx-3_::mCherry::tbb-2 3’UTR, P_rab-3_::mCherry::unc-54 3’UTR], nEx2958[P_ddx-23_::ddx-23::TagRFP-T::ddx-23 3’UTR, P_myo-2_::mCherry::unc-54 3’UTR], nEx2971[P_ddx-23_::ceDDX23::ddx-23 3’UTR, P_myo-2_::mCherry::unc-54 3’UTR #1], nEx2972[P_ddx-23_::ceDDX23::ddx-23 3’UTR, P_myo-2_::mCherry::unc-54 3’UTR #2], nEx2768[P_ceh-16_::DDX-23 cDNA::spGFPN::unc-54 3’UTR, P_ceh-16_::MAB-10 cDNA::spGFPC::unc-54 3’UTR, P_rab-3_::mCherry::unc-54 3’UTR], nEx2769[P_ceh-16_::DDX-23 cDNA::spGFPN::unc-54 3’UTR, P_ceh-16_::empty::spGFPC::unc-54 3’UTR, P_rab-3_::mCherry::unc-54 3’UTR], nEx2931[P_mab-10_::MAB-10::mCherry::mab-10 3’UTR, P_ceh-16_::DDX-23 cDNA::spGFPN::unc-54 3’UTR, P_ceh-16_::MAB-10 cDNA::spGFPC::unc-54 3’UTR, P_myo-2_::mCherry::unc-54 3’UTR], nEx2934[P_mab-10_::MAB-10::mCherry::mab-10 3’UTR, P_ceh-16_::DDX-23 cDNA::spGFPN::unc-54 3’UTR, P_ceh-16_::empty::spGFPC::unc-54 3’UTR, P_myo-2_::mCherry::unc-54 3’UTR]*

### Molecular cloning and transgenic strain construction

To endogenously tag the *mab-10* genomic locus at its C-terminus with GFP (transgene *n5909[mab-10::gfp])* using CRISPR-Cas9^73^, we generated a *mab-10::gfp* homologous repair template for Cas9-mediated *gfp* knock-in. We PCR-amplified a ∼3 kb genomic region centered on the *mab-10* stop codon from N2 genomic DNA and cloned the resulting fragment into the pCR-Blunt vector using the ZeroBlunt TOPO Cloning Kit (Thermo Fisher). Using the Gibson assembly kit (New England Biolabs), we inserted a GA-linker and *gfp* in-frame between the last amino acid codon of the *mab-10*gene and the stop codon, resulting in a homologous repair template with ∼1.5 kb homology arms. To avoid cleavage of the repair template by Cas9, we mutated the PAM motif of the Cas9 target site in the repair template. We also generated a Cas9-sgRNA plasmid to cleave upstream of the *mab-10* stop codon by inserting a targeting sequence (5’-GAAAAAGTAGCAATACAACA-3’) into the pDD162 Cas9-sgRNA plasmid^73^ by site-directed mutagenesis. Forward primer 5’-GAAAAAGTAGCAATACAACA GTTTTAGAGCTAGAAATAGCAAGT-3’ and reverse primer 5’-CAAGACATCTCGCAATAGG-3’ were used. We co-injected the homologous repair template, the *mab-10* Cas9-sgRNA plasmid, and an *unc-22* sgRNA co-CRISPR plasmid into wild-type N2 worms. Twitching progeny (indicative of successful *unc-22* co-CRISPR integration) were selected and genotyped for integration of *gfp* into the *mab-10* locus. Finally, the strain was backcrossed to wild-type N2 animals to remove the *unc-22* mutation.

To create the *ddx-23(n6026)* allele encoding DDX-23[I383F] and the *ddx-23(n6027)* allele encoding DDX-23[S365N] by CRISPR-Cas9, we used oligonucleotides as repair templates to introduce substitutions as previously described_74_. pDD162 was used as a template for site-directed mutagenesis to insert Cas9 targeting sequences for *ddx-23(n6026)*(5’-ACGACAAGAACATCGGGACT-3’) and *ddx-23(n6027)*(5’-AATGGAACGACAAGAACATC-3’).Temperature-sensitive *pha-1* co-conversion was performed as previously described^74^.

To generate the *ddx-23* rescuing transgene *nEx2848*, the region spanning 5 kb upstream of the *ddx-23* gene (*P_ddx-23_*), *ddx-23* genomic DNA, and the 3’UTR were PCR-amplified using forward primer 5’-GGTACAACCATGTTCAAATCAGCATCC-3’ and reverse primer 5’-CGTGAGTTTGGTTCTGGAGCTT-3’. This PCR fragment was cloned into a pUC19 vector and a Tag-RFP-T sequence was inserted immediately before the *ddx-23* stop codon to generate the *ddx-23* rescuing transgene *nEx2958. P_ddx-23_::ceDDX23::ddx-23 3’UTR (nEx2971* and *nEx2972)* was generated by HiFi DNA Assembly (New England Biolabs) of *P_ddx-23_*, wild-type human DDX23 cDNA codon-optimized for expression in *C. elegans* and containing 3 synthetic introns generated using gBlocks Gene Fragments (Integrated DNA Technologies), and the *ddx-23* 3’UTR in a pUC19 vector. For the split-GFP protein-protein *in vivo* interaction assays, all cloning was performed using the HiFi DNA Assembly system. The N-terminal GFP (spGFPN) plasmid was cloned by inserting the *ddx-23* cDNA and spGFPN fragment between a 2.5 kb *ceh-16* promoter and the *unc-54* 3’UTR. The C-terminal GFP (spGFPC) plasmid was cloned by inserting the *mab-10* cDNA (or an ATG start codon sequence for the “empty” control) and the spGFPC fragment between a 2.5 kb *ceh-16* promoter and the *unc-54* 3’UTR. Complete plasmid sequences of all plasmids are available from the authors upon request.

Transgenic strains were generated by germline transformation as described^75^. All transgenic constructs were injected at 1–50 ng/ml.

### Microscopy and image analysis

All *C. elegans* micrographs excluding electron micrographs are of lateral views of animals. Nomarski DIC and epifluorescence micrographs were obtained using a Zeiss Axio Imager Z2 compound microscope and Zeiss Zen software. Confocal microscopy was performed using the Zeiss LSM 800 instrument. The resulting images were prepared using ImageJ (National Institutes of Health) and Adobe Illustrator CS4 softwares. Image acquisition settings were calibrated to minimize the number of saturated pixels and were kept constant throughout each experiment. Graphs and indicated statistical analyses were performed using GraphPad Prism 6 software. For electron micrographs, animals were assayed by picking L4 males to plates, fixing 24 hours later and performing electron microscopic analysis as previously described.

### Fluorescence recovery after photobleaching (FRAP)

We immobilized one-day old adult animals with 10 mM levamisole (an acetylcholine agonist that induces muscle contraction) on 2% agarose pads for confocal imaging using the Zeiss LSM 800. We imaged hypodermal cells that were proximal to the cover slip. We identified two imaging regions of interest (ROIs) for each FRAP experiment (one for FRAP measurements, one for an imaging-associated bleaching control). We manually defined the FRAP ROI as the region around a given MAB-10::GFP focus. As a control, we performed FRAP studies of the diffuse MAB-10::GFP signal in hypodermal cells. To achieve complete bleaching of GFP in the FRAP ROI, bleaching was conducted with 25 rounds of 100% laser induction with a 488 nm laser. Hypodermal cells used to perform FRAP studies of diffuse MAB-10::GFP signals (“Diffuse”) were then tracked for at least 35 sec, while hypodermal cells used to perform FRAP studies of MAB-10::GFP foci (“Focus”) were tracked for at least 80 sec. Data were analyzed using ImageJ, measuring the fluorescence intensity for all frames and fitting the FRAP curve by an increasing exponential.

### Mutagenesis screen for enhancers of *mab-10* and mapping of *ddx-23*

To screen for enhancers of the *mab-10* seam-cell defect, we mutagenized *mab-10(tm2497)* mutants with ethyl methanesulfonate (EMS) as described previously^71^. The starting strain contained the *col-19::gfp* (*maIs105*) transgene, which shows a reduced expression level in *mab-10(lf)* adults relative to that of wild-type adults and served as a reporter for the heterochronic pathway. We used a Nikon SMZ18 fluorescence dissecting microscope to screen F2 progeny of mutagenized hermaphrodites for second-site mutations that further reduced the low hypodermal *col-19* expression levels of *mab-10(tm2497)* mutants. Screen isolates were backcrossed to an otherwise wild-type strain containing the *maIs105* reporter. *ddx-23(n5703)* and *ddx-23(n5705)* were identified by crossing the mutant isolate with a Hawaiian strain containing the *maIs105* reporter, isolating the F2 progeny with low hypodermal *col-19* expression and mapping the mutation using single-nucleotide polymorphisms^72^ and whole-genome sequencing. Transgenic rescue experiments demonstrated that these *ddx-23* mutations were the causative mutations, as described in the text.

### Seam cell count

The number of seam cells were quantified in one-day old adults, using either the *maIs105[col-19::gfp]* or *wIs78[ajm-1::gfp + scm::gfp]* transgenes. DIC and epifluorescence optics using a Zeiss 63x objective lens on a Zeiss Axio Imager Z2 compound microscope were used. The number of seam cells was scored by counting GFP-positive seam cell nuclei along the length of each animal.

### Yeast two-hybrid binding assay

*ddx-23* and *mab-10* cDNAs were cloned into pGADT7 and pGBKT7 plasmids, and the constructs were co-transformed into yeast strain Y2HGold (Takara). Co-transformants were selected on SD plates containing minimal supplements without leucine and tryptophan (SD/-Leu/-Trp) for two days at 30°C. To test for protein interaction, co-transformants were picked and grown overnight in SD/-Leu/-Trp broth. Cultures were diluted to a similar optical density at 600 nm, and 3 μl of each culture was spotted onto SD plates containing minimal supplements without tryptophan, leucine, histidine, and adenine (SD/-Leu/-Trp/-His/-Ade) and containing 10 mM of the competitive inhibitor of histidine synthesis 3-AT, and cultured for two days at 30 °C.

### Recombinant protein expression and protein purification

HiFi DNA Assembly into a pDEST17 (Thermo Fisher) backbone was used to create all bacterial expression vectors. For the recombinant DDX-23 expression vector, the *ddx-23* cDNA sequence followed by mNeonGreen and a 6xHis tag at its 3’ end was inserted into pDEST17. Site-directed mutagenesis of this plasmid was performed to create the DDX-23 I383F mutant and the DDX-23(ΔIDR) mutant (removing amino acid residues 2-265). For the recombinant MAB-10 expression vector, *mab-10* cDNA followed by mScarlet I and a 6xHis tag was inserted into pDEST17. Vectors expressing C-terminally 6xHis-tagged mNeonGreen and mScarlet I were created as “empty” controls.

For protein expression, plasmids were transformed into Shuffle T7 competent *E. coli* cells (New England Biolabs). A fresh bacterial colony was inoculated into LB medium containing ampicillin and grown overnight at 30°C. Cells were diluted 1:100 in 1 liter of LB medium containing ampicillin and grown at 30°C until the OD600 reached between 0.5 and 0.6. IPTG was added to 0.5 mM, followed by incubation at 16°C for 18 hr. Cells were collected and stored frozen at −80°C. For protein purification, bacterial pellets were resuspended in 15 ml of lysis buffer (20 mM Tris-HCl pH 7.4, 500 mM NaCl, 10 mM imidazole, 14 mM β-mercaptoethanol (ME), 10% (v/v) glycerol, 1% Triton-X) with Complete EDTA-free protease inhibitors (Roche) and 15 mg of lysozyme, and sonicated. The lysates were cleared by centrifugation at 10,000 x*g* for 30 min and added to 1 ml of Ni-NTA agarose (Qiagen) pre-equilibrated with the same lysis buffer, and rotated at 4°C for 2 hr. The slurry was centrifuged at 3,000 *xg* for 3 min. The resin pellets were washed 5 times with 4 ml of wash buffer (20 mM Tris-HCl pH 7.4, 500 mM NaCl, 14 mM β-ME, 10 % (v/v) glycerol, and 25 mM imidazole), followed by centrifugation as above. Protein was eluted twice for 15 min with 2 ml of Ni-NTA Elution Buffer (20 mM Tris pH 7.4, 500 mM NaCl, 14 mM β-ME, 10% (v/v) glycerol, and 250 mM imidazole). We further purified the DDX-23 and MAB-10 proteins using heparin columns. For DDX-23-mNeonGreen-6xHis and DDX-23[I383F]-mNeonGreen-6xHis, eluates were pooled and diluted in 20 ml of 1:1 mixture of Heparin Binding Buffer (20 mM Tris pH 7.4, 50 mM NaCl, 1% (v/v) glycerol, 2 mM DTT) and Heparin Elution Buffer (20 mM Tris, pH 7.4, 1M NaCl, 1% (v/v) glycerol, 2 mM DTT). For DDX-23(ΔIDR)-mNeonGreen-6xHis and MAB-10-mScarlet I-6xHis, eluates were diluted in 20 ml of Heparin Binding Buffer. The protein sample was applied to a pre-equilibrated heparin column with a flow-rate of 1 ml/min, followed by a column wash using the same buffer in which the protein had been diluted. DDX-23-mNeonGreen-6xHis and DDX-23[I383F]-mNeonGreen-6xHis proteins were eluted in Heparin Elution Buffer; DDX-23(ΔIDR)-mNeonGreen-6xHis protein was eluted in 7:3 ratio of Heparin Binding Buffer : Heparin Elution Buffer; MAB-10 protein was eluted with a series of step elutions with Heparin Binding Buffer : Heparin Elution Buffer ratios ranging from 3:1 to 1:1. All recombinant mNeonGreen or mScarlet I fusion proteins were concentrated using Vivaspin 500 Centrifugal Concentrators (10,000 MWCO). We analyzed the proteins using a 12% acrylamide gel followed by Coomassie staining to assess protein purity. Protein concentration was measured using the Pierce 660nm Protein Assay Kit (Thermo Fisher).

### *in vitro* protein interaction assay

As described in the figure legends, recombinant proteins were added to solutions at varying concentrations with 200-220 mM final NaCl with or without 10% PEG-8000 as crowding agent in the reaction buffer (50 mM Tris-HCl pH 7.4, 10% glycerol, 1 mM DTT). The protein solution was loaded onto glass slides within a silicone isolator (Sigma) and imaged using a Zeiss LSM 800. For all experiments at least three independent mixtures were analyzed. For better visualization, micrographs were color-inverted using ImageJ.

### Gene Expression Analyses

For RNA-Seq experiments, total RNA from three biological replicates of age-synchronized young adult hermaphrodites (200 in total, picked manually) of wild-type, *lin-29(n836), mab-10(tm2497), and ddx-23(n5705)* genotypes were prepared using a BeadBug microtube homogenizer (Sigma) and 0.5 mm zirconium beads (Sigma). RNA was extracted using the RNeasy Mini kit (QIAGEN) according to the manufacturer’s instructions. RNA-Seq was performed using the Illumina NextSeq500 platform, and data were analyzed using standard protocols^76^. An adjusted p-value of < 0.05 was used to identify differentially expressed genes between the wild type and mutants. There were 149 genes -- including larval-specific collagen genes and several *hedgehog*-related (*grd)* genes and putative *hedgehog* receptor (*ptr*) genes -- that were up-regulated in all three mutants (*lin-29, mab-10,*and *ddx-23* mutants) relative to wild-type animals.

### Statistical Analysis

Unpaired t-tests were used to compare half-maximal recovery times from FRAP studies of MAB-10::GFP in foci and diffuse signals, numbers of seam cells that undergo extra seam cell division between strains, normalized gene expression abundance log_2_(fpkm+1) of multiple genes between strains, the number of homotypic droplets of control mNeonGreen alone to that of purified DDX-23- mNeonGreen, the number of homotypic droplets of control mScarlet I alone to that of purified MAB-10-mScarlet I, and the number of MAB-10 droplets in mixed heterotypic assays with control, wild-type DDX-23, mutant DDX-23[I383F] or DDX-23(ΔIDR) proteins. Fisher’s exact test was used to compare the number of animals that have MAB-10::GFP foci in adult hypodermal nuclei in wild-type versus *ddx-23(n5703)* or *ddx-23(n5705)* mutants. Statistical tests were performed using GraphPad Prism 6 software. Biological replicates were performed using separate populations of animals.

### Accession numbers

The GEO accession number for the RNA-Seq dataset in this paper is GSE151399.

## Acknowledgments

We thank V. Ambros and S. V. Lee for reagents; the *Caenorhabditis* Genetic Center, which is funded by the NIH Office of Research Infrastructure Programs (P40 OD010440) and the National BioResource project for strains; and V. Dwivedi, D. Ghosh, A. Corrionero, K. Burkhart, and J. Kong for comments concerning the manuscript. This work was supported by NIH grant R01GM024663. A.D. was supported by the Uehara Memorial Foundation and Japan Society for the Promotion of Science postdoctoral fellowships and by the Howard Hughes Medical Institute. H.R.H. is an Investigator of the Howard Hughes Medical Institute and the David H. Koch Professor of Biology at MIT.

## Author Contributions

H.R.H. supervised the project. A.D. initiated the project and designed experiments. A.D., G.D.S. and R.D. performed the experiments. All authors contributed to data analysis, interpretation, and manuscript preparation.

## Declaration of Interest

The authors declare no competing interests.

## Notes

### Competing Interest Statement

The authors have declared no competing interest.

